# Effectiveness of Demand creation promotions and demand creation personnel in creating demand for Voluntary Medical Male Circumcision in Chitungwiza district, Zimbabwe in 2016

**DOI:** 10.1101/331397

**Authors:** Taurai Matikiti, Tsitsi P Juru, Notion Gombe, Peter Nsubuga, Mufuta Tshimanga

**Affiliations:** Zimbabwe Community Health Intervention Research Project (ZiCHIRE); Department of Community Medicine, University of Zimbabwe; Global Public Health Solutions, USA

**Keywords:** VMMC, demand creation promotions, demand creation personnel, service providers, VMMC clients, effectiveness

## Abstract

Zimbabwe is one of the 14 countries in eastern and southern Africa that have adopted Voluntary Medical Male Circumcision (VMMC) as an HIV prevention intervention in response to WHO’s recommendation for countries with generalised high HIV epidemics and low VMMC prevalence. However, since 2013 when VMMC was scalled up in Zimbabwe, there was a general low uptake of the VMMC programme particularly on the target age group 20-29 years which has an immediate reduction in the HIV burden. The failure of VMMC uptake in the priority age group prompted the need to analyse the effectiveness of demand creation promotions and personnel used in creating demand for VMMC in Chitungwiza district, Zimbabwe. We employed judgmental sampling, a non-probability sampling technique where we interviewed VMMC clients (n=50) and service providers (n=10) using self administered questions, and community mobilisers (n=10) and demand creation teams (n=3) using face-to-face interviews based on their experience, knowledge and professional judgment. We also randomly analysed client records in the form of 50 Client Intake Forms (CIF) books. We found out that Community mobilisers and Demand creation officers were effective in mobilising clients in the age group 10-15 years and 16-30 years respectively. The use of clinicians (nurses) was also found to be useful in creating demand for VMMC. We also found out that intensifying campaigns during school holidays, the use of tent-based/caravan campaigns and the door to door campaigns were most effective strategies under demand creation promotions. We concluded that there is need to increase demand creation officers and qualified community mobilisers. To regulary train and motivate current community mobilisers as well as increasing the use of clinicians(nurses) in demand creation. We recommended the need to increase the number of mobile caravans and intensifying on the door to door campaigns in the district.

## INTRODUCTION

Voluntary Medical Male Circumcision(VMMC) is the removal of the fold of the skin that covers the head of the penis by a medically trained service provider on a client who has gone through the counselling process and consent to undergo the procedure(1–3). WHO and UNAIDS recommended VMMC as an additional strategy for HIV prevention after several types of research in Kenya, Uganda and South Africa showed that VMMC reduces the risk of female-to-male transmission of HIV by approximately 60%(4,5). Countries with high HIV prevalence and low levels of male circumcision were targeted resulting in 14 countries in eastern and southern Africa being prioritised,and Zimbabwe was one of the countries(3–5).

In Zimbabwe, VMMC was adopted by the Ministry of Health and Child Care (MOHCC) as part of a comprehensive HIV prevention package in 2007,and the scale-up phase started in 2013 after some pilot studies were carried out and partners were secured to fund the programme(1). Population Services International Zimbabwe, International Training and Education Centre for Health, Zimbabwe Association of Church-related Hospital, Integrated Support Programme and Zimbabwe Community Health Intervention Research Project (ZiCHIRe) were partners supporting MOHCC in implementing VMMC in various districts of Zimbabwe(6).ZiCHIRe offered support in six districts including Chitungwiza district.

Chitungwiza district is in Mashonaland East province of Zimbabwe and has four static sites that provide VMMC and has a mobile caravan where VMMC services are also offered. MOHCC came up with a target to circumcise 1.3million males aged above 10 years and to circumcise 80% of HIV negative men aged between 13-29 years by 2017. Particular emphasis was on the 20-29 year age group to have an immediate reduction in the HIV burden(6). By year-end 2015 the 10-15 year age group contributed more than 50% of the total number circumcised across all districts in Zimbabwe(6).MOHCC employed various strategies such as advertising campaigns, public relation campaigns, market segmentation, demand creation promotions among others, and its partners to meet the targets(1). However, despite these strategies, the annual target for VMMC has not been achieved. The failure of VMMC uptake in the priority age group prompted the need to analyse the effectiveness of demand creation promotions and personnel used by ZiCHIRe in creating demand for VMMC. Chitungwiza district was considered for the study because the district had not achieved its target, most demand creation promotions were carried out in it, it had significant cadres in demand creation team and it was the nearest district for the researchers who were based in Harare.

Demand creation promotions consist of mostly short-term incentives designed to motivate clients to adopt VMMC. In service marketing, when applied promotions will motivate consumers to purchase a product immediately and in larger quantities(7,8). It is offering over and above what would normally be provided to the customer (9). In VMMC when used effectively, promotions can create quick demand and persuade the resistant clients to consider VMMC. Demand creation campaigns include such activities as the door to door campaigns(10), intensive VMMC holiday campaigns (11), tent-based/caravan campaigns, short message service (SMS) campaigns(12), Smartphone raffles(13) and offering of free men’s’ health checkups. In this study,we evaluated the effectiveness of demand creation promotions used in Chitungwiza district.

Demand creation personnel are the people directly employed and used to create demand for VMMC by communicating with prospective clients and other stakeholders (7). To evaluate the effectiveness of demand creation personnel, data from demand creation officers, community mobilisers, male champions, demand creation field assistants and barbers were analysed. Interpersonal communication through community mobilisers was noted as the most effective strategy as it catalyses action(11,14,15).

This study aimed to evaluate the effectiveness of demand creation promotions like the door to door campaigns and demand creation personnel like community mobilisers and demand creation officers used in creating demand for VMMC. This was to help in recommending channelling of resources to activities and personnel that will result in an increased update of VMMC in the priority age group to have an immediate impact on the HIV burden. The findings of the study would also help ZiCHIRe and ultimately MOHCC in crafting evidence-based strategies that create demand for VMMC programme

## METHODS

### Study design

We used descriptive research design in conducting a study on the effectiveness of demand creation promotions and demand creation personnel used in creating demand for VMMC using a quantitative approach.

### Study settings

We conducted the study in Chitungwiza district in the period July to September 2016. All the four static sites in the districts that offered VMMC –Zengeza 3 Clinic, Seke South Clinic, Seke North Clinic and Chitungwiza Central Hospital and a mobile caravan were considered in carrying out this study with particular reference on VMMC clients, community mobilisers, service providers and demand creation officers from Chitungwiza districts. Chitungwiza is mainly urban, but it also has part of a rural setting. It has many council-run schools,and private schools and its population is shared between formal and informal employment with a significant number of its people working in the capital city –Harare.

### Study population

We conducted the study among VMMC clients, community mobilisers, VMMC service providers and demand creation officers. VMMC clients comprised prospective clients coming for VMMC services and circumcised clients coming for reviews.

### Sample size

We recruited the whole population of critical stakeholders in VMMC programme. Our sample consisted of 73 participants; comprising 25 VMMC circumcised clients who were coming for reviews or we followed up, 25 VMMC prospective clients coming for the service, three demand creation officers who were operating in Chitungwiza on a rotational basis, 10 VMMC service providers and 10 VMMC community mobilisers. The sample was based on the number of VMMC service providers in the district, community mobilisers and weekly average of 100 circumcisions. However, 10% of VMMC clients did not submit their questionnaires,and the interviewed total number became 68.

### Sampling

We also randomly sampled 50 Client Intake Books (CIF) to find out what was recorded as the source of VMMC information by clients who come for VMMC services. There was even distribution of age for the fifty clients chosen with aspecific emphasis on 20-29 age groups. The sample was considered to represent a comprehensive picture of the views of the total population because age, experience and knowledge in VMMC were considered in the determination of theeffectiveness of VMMC promotional strategies.

### Sampling procedure

We employed judgmental sampling, a non-probability sampling technique where we selected VMMC clients, community mobilisers, demand creation teams and service providers based on their experience, knowledge and professional judgment. We also randomly choose and analysed client records in the form of 50 Client Intake Forms (CIF) books.

### Data Collection Procedures

We used self-administered questionnaires on10 service providers and 50 clients while face-to-face interview guided by a questionnaire was used on community mobilisers and demand creation officers. We conducted a desk review of 50 CIF books and client register to determine what clients stated as the source of information to consider VMMC. Information from client register was also used to verify information on CIF particularly the age of clients and to follow up on circumcised clients to be interviewed

We physically gave the research questionnaires to VMMC clients and service providers. On occasions where we were not present at some health facilities, we appointed one service provider to give the questionnaires to clients and guide them in answering the questionnaire. We apprised one service delivery member on all the facilities offering VMMC in Chitungwiza on the objectives of the study to make them guide clients in completing the questionnaire. Interviews with demand creation officers and community mobilisers were carried out at their convenient times.

### Data Analysis

We analysed open-ended questions and subsequently used excel spreadsheet to compute data frequencies. Tables, bar graphs and pie charts were used to present the data. On-demand creation promotions, we focused on five elements which were door to door campaigns, school holiday campaigns, tent-based/caravan campaigns and offering of free men’s health check-ups. These elements were analysed to see if they were useful in creating demand for VMMC by summing up the ratings by respondents in percentages. Respondents were asked to rate the effectiveness of the above elements of demand creation promotions and were also given the opportunity to suggest other options. Effectivenesswas rated on a five-pointLikert scale as follows; not effective, not sure, slightly effective, moderately effective and very effective.

We analysed data from community mobilisers, demand creation officers, demand creation field assistants, male champions and use of barber as mobilisers under demand creation personnel and participants were also given the opportunity to provide other suggestions. We asked all respondents to state the demand creation personnel they perceive to be effective in different age groups 10-15 years, 16-29 years and over 30 years as per MOHCC categories. We also asked them to rate in general the effectiveness of various demand creation promotions in creating demand for VMMC. Effectivenesswas rated on a five-pointLikert scale as follows; not effective, not sure, slightly effective, moderately effective and very effective. Data from CIF were collected and presented on a pie chart.

### Ethical Considerations

In this study, ethical consideration was given to avoid violating the respondents’ rights to privacy. Permission to conduct the study was obtained from ZiCHIRe authorities. The information provided by the respondents was treated with the confidentiality and was only usedfor the study.

## RESULTS

### Demographic Distribution of VMMC Clients

We collectedinformation on thedemographic distribution of VMMC clients which include age, educational level, occupational status and the marital status from 45 clients after 5 clients did not submit their questionnaire. The results are presented in Table1.

**Table 1:** Demographic distribution of VMMC clients interviewed in Chitungwiza district in August 2016.

| **Variable** | **Category** | **Number** | **Percentage (%)** |
| --- | --- | --- | --- |
| Age Distribution | 10-15 years | 5 | 11 |
|  | 16-29 years | 34 | 76 |
|  | Over 30years | 6 | 13 |
| Educational level<br>(Highest Level) | Primary | 1 | 2 |
|  | Secondary | 10 | 23 |
|  | Certificate | 23 | 51 |
|  | Degree | 9 | 20 |
|  | Post graduate | 2 | 4 |
| Marital Status | Single | 25 | 56 |
|  | Married | 16 | 36 |
|  | Divorced | 2 | 4 |
|  | Widowed | 2 | 4 |
| Occupation | Self employed | 21 | 47 |
|  | Employed | 11 | 24 |
|  | Not working | 13 | 29 |

The majority of VMMC clients (76%) were between 16 and 29 years of age. The majority of VMMC clients (51%) had attained certificates in various programmes, only 2% were educated to primary level, 20% had degrees, and 23% had achieveda secondary education.

Of the clients selected 56% (n=45) of them were single,and 36%(n=45) were married. On occupation the majority of VMMC clients (47%) interviewed were self-employed, 29% unemployed and only 24% were employed.

### Educational level and years of experience in VMMC of Chitungwiza community mobilisers sampled

We collected information on theeducationalleveland years of experience of community mobilisers in Chitungwizaand the results were presentedin Table 2.

**Table 2:** Chitungwiza Community mobilisers’ educational level and years of experience in VMMC, August 2016 Educational Level.

| **Highest Qualification** | **Frequency** | **Response Rate (%)** |
| --- | --- | --- |
| Primary | 1 | 10 |
| Secondary | 7 | 70 |
| Certificates | 2 | 20 |
| **Total** | **10** | **100** |

**Years of experience in VMMC**
| **Number of Years** | **Frequency** | **Response Rate(%)** |
| --- | --- | --- |
| 3 | 2 | 20% |
| 4 | 5 | 50% |
| More than 5 | 3 | 30% |

The majority of community mobilisers 70% (n=10) attained their secondary education, and all the community mobilisers had more than three years of experience in VMMC.

### Clients’ Source of information about VMMC

Information about how clients learnt about VMMC services from 50 Client Intake Form books were collected,and Figure 1 shows the results.

**Figure 1:**
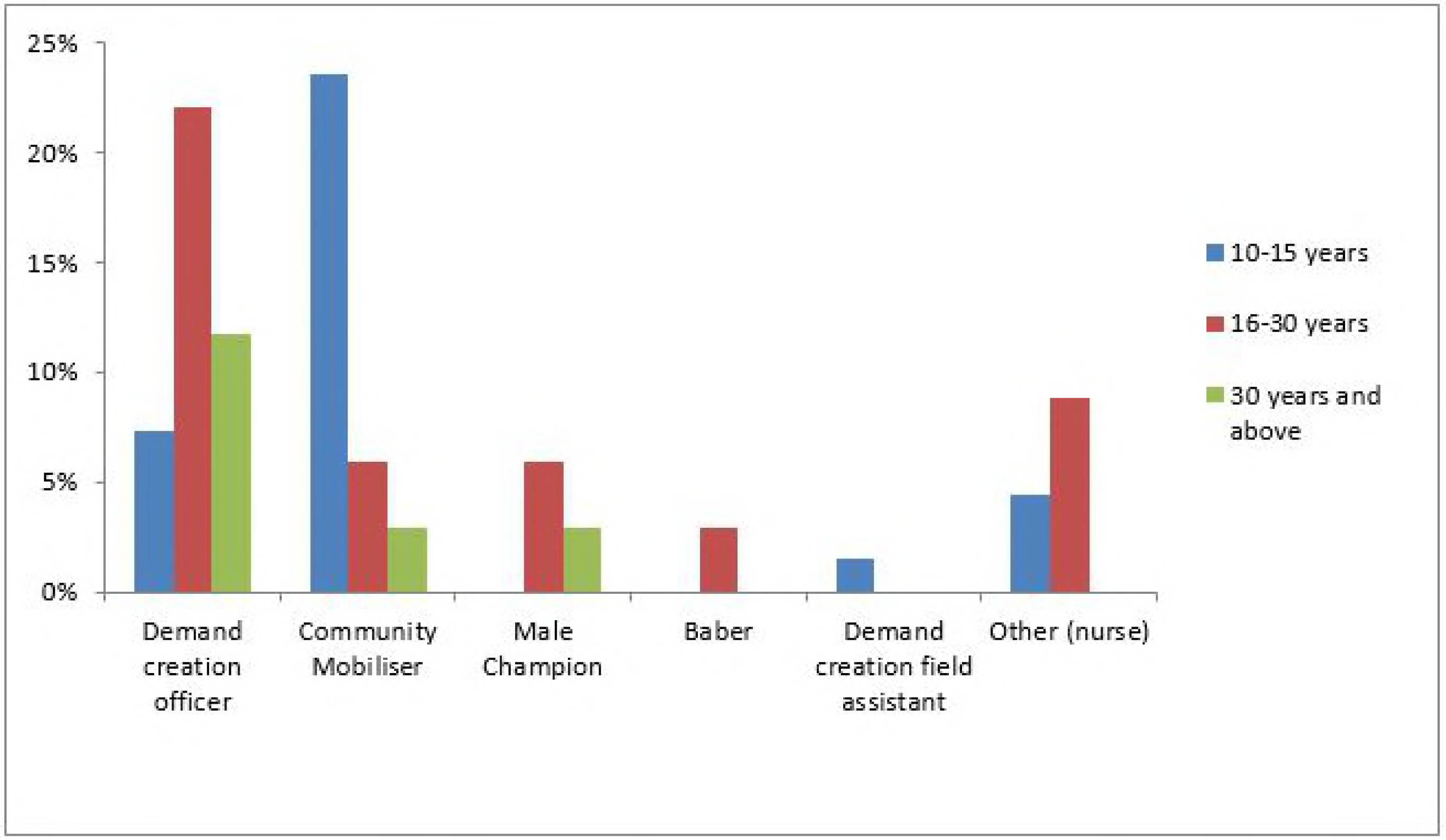
Chitungwiza VMMC Clients source of information from CIF, Aug 2016.

Information from CIF shows that most clients mentioned community mobilisers as the source of VMMC information, with radio 16%, television 12% and Health worker cited as the other common source of information in Chitungwiza district.

### The effectiveness of Demand creation personnel in different age groups

Information from questionnaires and interviews carried out with participants on the effectiveness of demand creation personnel in different age groups were collected and presented in Figure 2.

**Figure 2:**
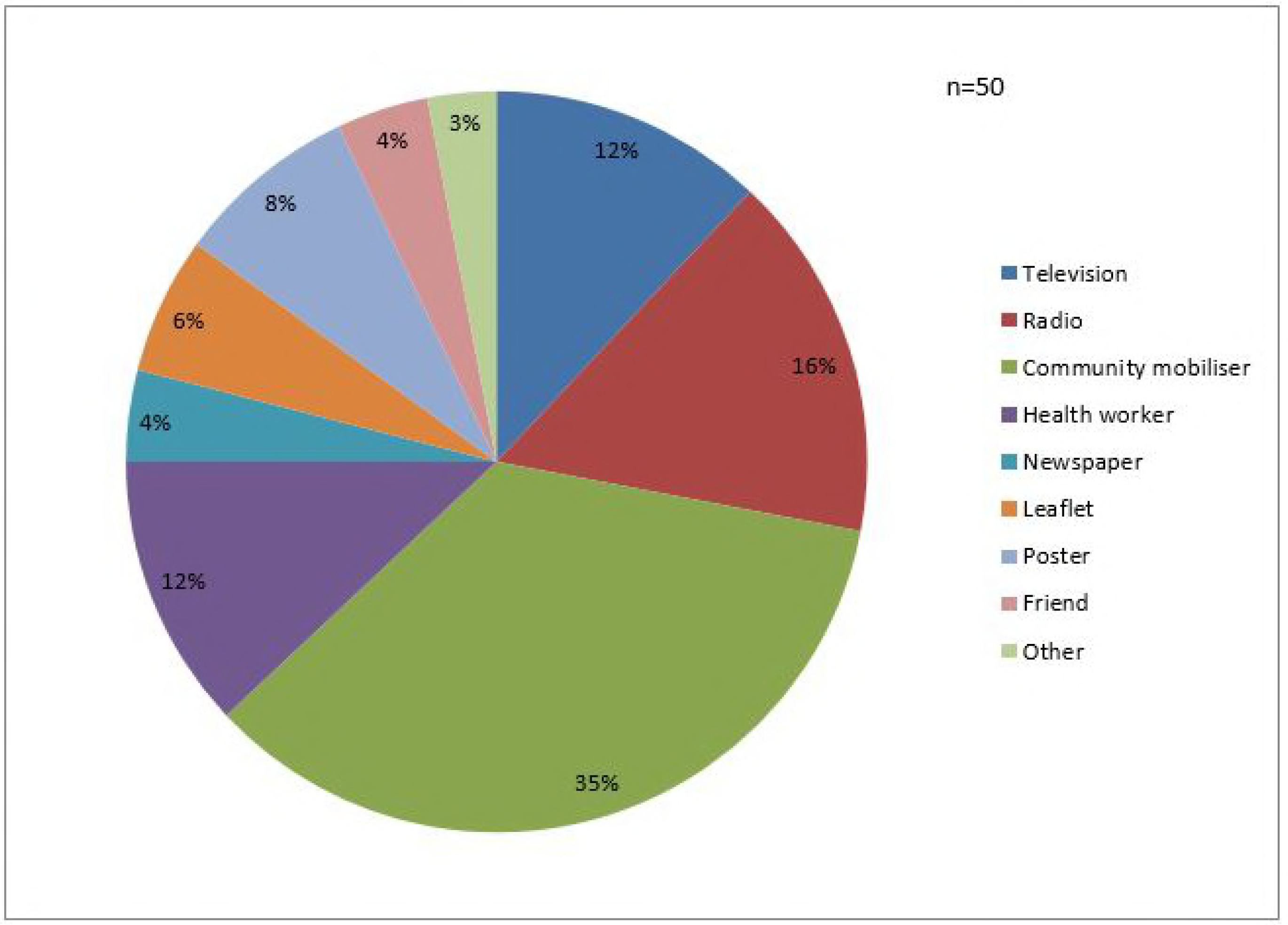
Effectiveness of Chitungwiza demand creation personnel in different age groups, Aug 2016.

Demand creation officers (23%) and Community mobilisers (24%) were effective in mobilising clients in theage group 16-30 years and 10-15 years respectively. Participantsadded nurses,andthey were next after demand creation officers and community mobilisers in creating demand for the 10-30 years age group.

### The effectiveness of VMMC demand creation personnel in creating demand for VMMC

In rating demand creation personnel in creating demand for VMMC services, all participants-community mobilisers, demand creation officers, VMMC clients and service providers were asked to rate their view,and their results were presented in Table 3.

**Table 3:**
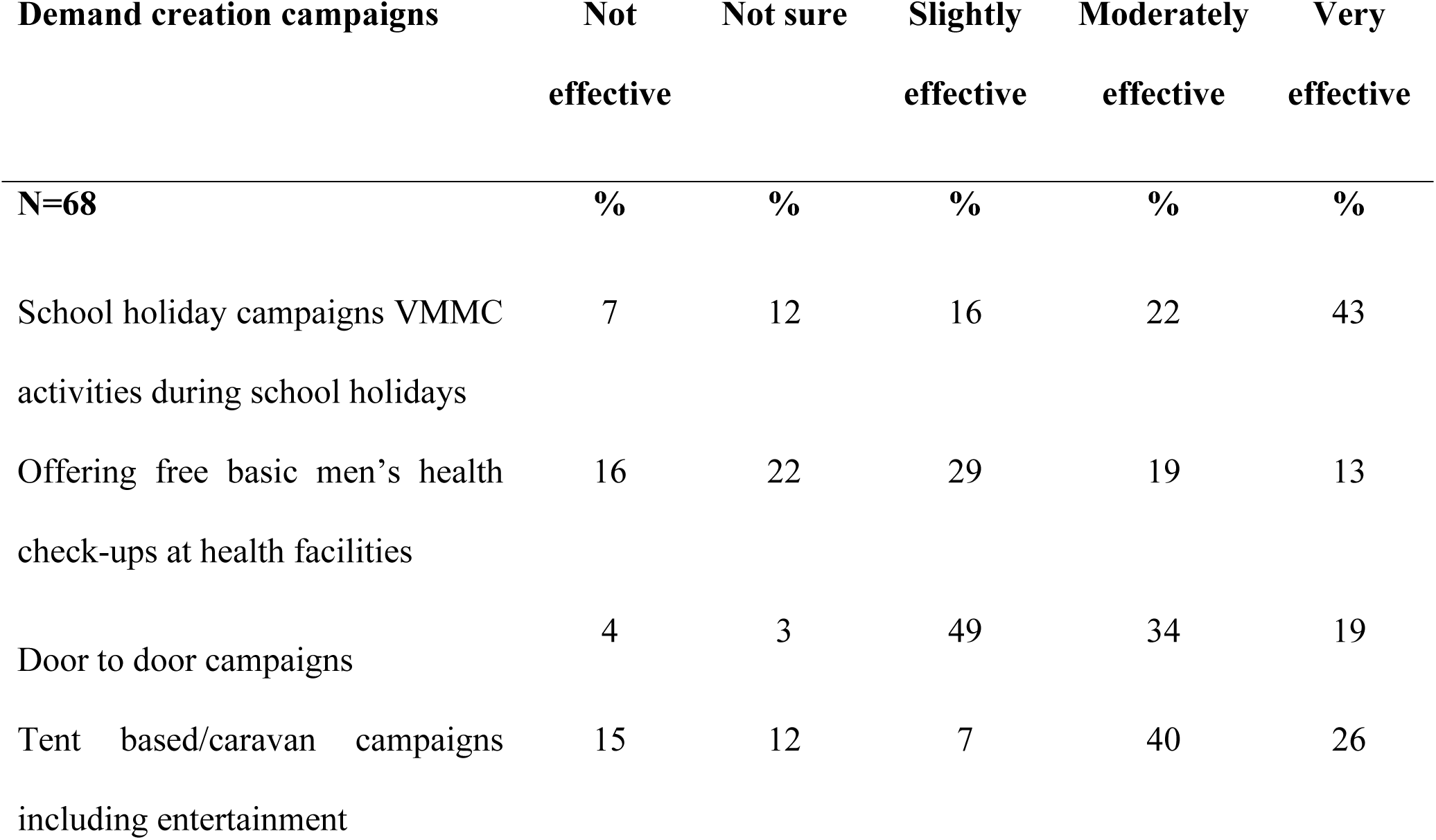
ZiCHIRe demand creation campaigns used in Chitungwiza district.

Results from the research showthat community mobilisers (44%), demand creation officers (49%) and the use of male champions (30%) were veryeffective in persuading boys or men to undergo VMMC. Forty percent of the respondents indicated that a barbershop mobiliser model is not useful in creating demand for VMMC services. The use of nurses (29%) and mobile caravan manned with service delivery and demand creation team (28%) were noted to be very effective and effective respectively.

### The effectiveness of demand creation promotionsin creating demand for VMMC in Chitungwiza

All participants(VMMC clients, service providers, demand creation teams and community mobilisers) were asked to rate their views on demand creation promotions such as tent based campaigns, door to door campaigns among others, and Table 4 shows the responses.

**Table 4:**
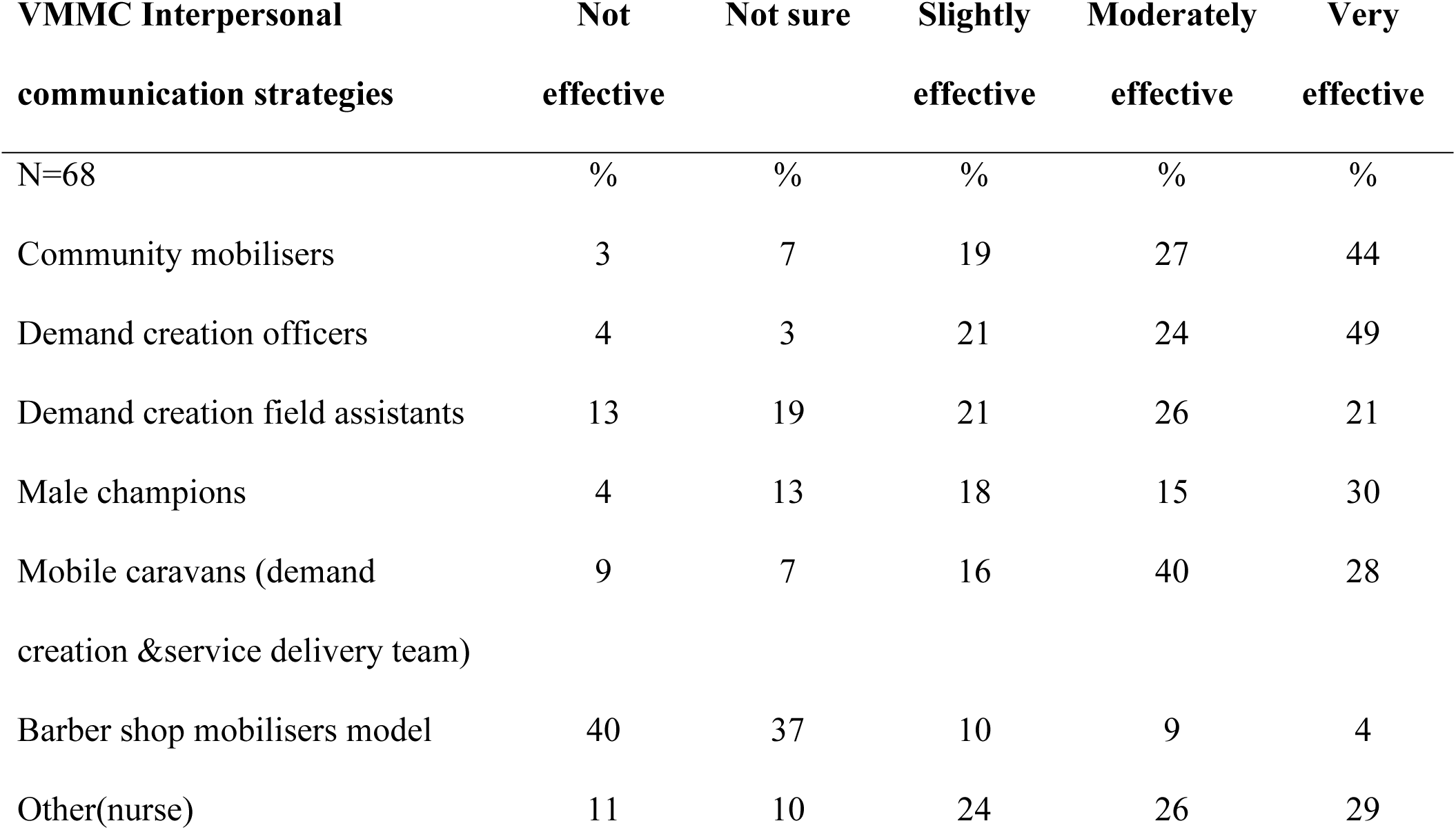
ZiCHIRe VMMC interpersonal communication strategies in Chitungwiza district in August 2016.

| **VMMC Interpersonal**<br>**communication strategies** | **Not**<br>**effective** | **Not sure** | **Slightly**<br>**effective** | **Moderately**<br>**effective** | **Very**<br>**effective** |
| --- | --- | --- | --- | --- | --- |
| N=68 | % | % | % | % | % |
| Community mobilisers | 3 | 7 | 19 | 27 | 44 |
| Demand creation officers | 4 | 3 | 21 | 24 | 49 |
| Demand creation field assistants | 13 | 19 | 21 | 26 | 21 |
| Male champions | 4 | 13 | 18 | 15 | 30 |
| Mobile caravans (demand<br>creation & service delivery team) | 9 | 7 | 16 | 40 | 28 |
| Barber shop mobilisers model | 40 | 37 | 10 | 9 | 4 |
| Other(nurse) | 11 | 10 | 24 | 26 | 29 |

In exploring promotions that would stimulate immediate or quick demand for VMMC, school holiday campaigns or intensive VMMC activities during school holidays (43%), door to door campaigns (19%) and tent based campaigns (26%) were cited to be very effective. Free basic men’s health check-ups at health facilities can act as ahook to persuade men who come for the service to adopt VMMC and were cited as slightly effective (29%).

## DISCUSSION

In our evaluation of the effectiveness of personnel used in demand creation and demand creation promotions, we found out that demand creation officers and community mobilisers were the most effective cadres in creating demand for VMMC in the target age group of 16-29 years in Chitungwiza district. The target age group is the sexuallygroup active and has the highest incidence of new HIV infections and is the age that MOHCC is emphasising on to have an immediate impact on HIV reduction(1,16). We also found out that intensifying campaigns during school holidays and the use of tent-based/caravan campaigns were the most effective strategies that created demand for VMMC under sales promotions.

Community mobilisers and Demand creation officers were effective in mobilising clients in theage group 10-15 years and 16-30 years respectively. The reason why demand creation officers were effective in 16-30 years we attributed to the fact that the majority of VMMC clients were educated with 51% of them having attained certificates in various programmes and it would require intellectual cadres to motivate educated clients to consider VMMC. All the demand creation officers had experience in working with the communities, and they attended various MOHCC workshops which makes them acquire adequate knowledge on demand creation.However, since the majority of community mobilisers (70%) only attained to thesecondarylevel, they were overpowered by demand creation officers in mobilising clients in the category 15-30 years. They, however, used their adequate experience and knowledge to create demand in the lower age group (10-15 years). The three demand creation officers had degrees,and their intellectual capacity helped them to create demand in the target age group.

Overall community mobilisers were leading in creating demand for VMMC in communities(17). Information from client records-CIF books also confirms community mobilisersas the primarysource of VMMC information. In an earlier study carried out in Tanzania,Mahleret al. (2011) also recommended the use of community mobilisers in helping create demand for VMMC (11). The same community mobilisers were noted bySgaieret al. (2014) in their study when they recognised interpersonal communication through community mobilisers in addressing individual concerns as one of the most effective strategies in creating demand for VMMC(14).

The use of barber as mobiliser was not found to be ineffective in creating demand for VMMC. Male champions were found to be effective,but they were not used in Chitungwiza. The respondents were pointing to the celebrities like Jah Prayzah & Sulu-the local artists used on some VMMC posters and billboards to actively mobiliser clients for VMMC. The use of their wives was also suggested as useful in creating demand for VMMC. The use women in convinced men to undergo VMMC were recommendedin earlier studies by Cook et al. (2015) in Zambia and Brito et al. (2009) in the Dominican Republic(18,19).However, this was not implemented in Chitungwiza district, and further analysis after its implementation will be required to substantiate theeffectiveness of male champions in demand creation.

The use of clinicians (nurses) was found to be useful in creating demand for VMMC. Despite the fact that we did not include nurses in our evaluation of demand creation personnel, they were mentioned by respondents under other options,and this could point to the fact that they are very effective in creating demand for VMMC. Nurses were next after demand creation officers and community mobilisers in creating demand for the 10-30 years age group. School holiday campaigns or intensive VMMC activities during school holidays were noted to be valuable if well planned and adequately resourced. This finding was consistent with results from a study carried out by Ashengoet al. (2014) on VMMC in Tanzania and Zimbabwe and noted that in Zimbabwe campaigns were of use in the age group 10-24 years were accessed during a period of campaigns(20). The use of intensive campaign like school holiday campaigns can have great impact in generating high volume demand for VMMC provided that enough human resources and commodities are in place to ensure that quality standards are maintained(11).

We found out that tent-based/caravan campaigns were useful as the entertainment will help to gather people together and once people are gathered it will be easy to promote VMMC services. Adequate human resources and commodities must be in place to ensure service is offered to clients who opt for circumcision efficiently. Tent based campaigns approach can be helpful in generating high volume demand for VMMC(11).

The door to door campaigns and offering freebasic men’s health check-ups at health facilities were noted to have potential to create demand for VMMC,butthe strategies have not often been used in Chitungwiza to be measured objectively. The door to door campaigns have not been done much in Zimbabwe and as such have not yielded more results compared to other methods(10). However if adequately planned and resourced the strategies have thepotential to generate numbers. The offering of free basic men’s health checkupswasdone in 2015,and it has potential to act as a hook on men who have visited the facility to adopt VMMC. Service providers will have the opportunity to recommend VMMC. Howeverthis should be adequately advertised, planned and adequatelyresourced to be successful.

Our study had thelimitation that we were only confined to one district because of the time frame for the project. However to mitigate against the limitationconsider clients from the rural and urban setup of Chitungwiza district. We also selected community mobilisers, demand creation teams and service providers based on their experience, knowledge and professional judgment. We also used client records in analysing data.

## Conclusion

ZiCHIRe should increase the number of mobile caravans in the district such that at each shopping centre there is a mobile caravan. These are more appealing in campaigns and have potential in creating demand for VMMC. The door to door campaigns should also be intensified as they are effective in creating demand foor VMMC.

We concluded that there is need to increase demand creation officers who were found to be effective in the priority age group. There is need to increase the use of qualified community mobilisers who can address well in secondary schools and tertiary institutions to improve their effectiveness. Regular training and motivating current community mobilisers would also increase their effectiveness as most clients in the age group 16-29 years were found to be educated. Increased use of nurses in demand creation was found to be effective as well.

## Recommendation

We recommended full implementation of caravan/tent based campaigns, the door to door campaigns, the offering of free basic men’s health checks ups and continue intensifying campaigns during school holidays. There should be adequate planning and resourcing with human resources, Information, Education and Communication (IEC) materials, promotional materials and commodities of these promotion strategies to yield more numbers.

On-demand creation personnel we recommended increasing the number of demand creation officers in the district, recruiting educated community mobilisers and increase on refresher training on the current community mobilisers as well as motivating them by increasing on incentives. We also recommended increased use of nurses in demand creation for VMMC services.

